# Cell-Based Potency Assay for Anti-CD3-Anti-CD19 Diabody

**DOI:** 10.1101/2025.04.15.648836

**Authors:** Tania L. Weiss, Justine Paniagua, Tonny Johnson, Timothy P. Cripe, Peter Ralph

## Abstract

A cytotoxicity assay was developed to measure the potency of GP101, a single chain diabody. GP101 is comprised of two linked Fv domains, with one that binds CD3 (expressed on T cells), and the other that binds CD19 (expressed on B cells). GP101 directs CD3 positive T lymphocytes to CD19 positive B lymphocytes. This immunotherapy redirects CD3+ T cells to CD19-expressing B-cell malignancies, enabling T-cell-mediated tumor cell lysis. We developed a GMP cell-based potency assay to satisfy the FDA requirements. The potency assay uses a cytotoxic T cell line, and a B cell lymphoma cell line as the target. The target lymphoma B cell line is prelabeled with a fluorescent dye, and upon T cell mediated killing, the fluorescence dye is released and detected using a fluorometer. The emitted fluorescence is proportional to the dose of GP101. Greater than 90% of the target B cells were killed within two hours exposure *in vitro* with the lowest amount of detectable killing at 60pg/mL GP101. The assay is suitable for measuring purified GP101, or GP101 expressed by cells transduced by GP101 plasmid or AAV preparations, and bioactivity in animal or human blood. This novel assay met GMP/GLP compliance, allowing a quantifiable and reproducible measure of efficacy, ensuring batch-to-batch consistency, and met safety and effectiveness regulatory requirements. This potency assay may be applicable for testing other CD3-CD19 T cell engagers and suitable for developing other diabody mediated potency assays with appropriate antigens.

## 1. Introduction

A diabody, constructed from the variable regions of two different antibodies, co-engages two cell types expressing different antigens (1, 2). Blinatumomab, marketed as Blincyto®, is a diabody and the first bispecific T-cell engager (BiTE^TM^) to receive FDA approval, which was granted in 2014 for the treatment of relapsed or refractory B-cell precursor acute lymphoblastic leukemia (ALL). Blinatumomab functions by simultaneously binding to a CD19 epitope on B cells and a CD3epsilon epitope on T cells, facilitating T-cell-mediated lysis of normal or malignant B cells. Its clinical efficacy has been demonstrated in both relapsed/refractory and minimal residual disease-positive ALL, significantly improving overall survival rates (3-8).

Nationwide Children’s Hospital (Ohio, USA) has developed an Adeno-Associated Virus (AAV) gene therapy to deliver the gene encoding GP101 (9) similar to blinatumomab to treat B cell leukemias and lymphomas. This AAV therapy, administered as a single off-the-shelf dose, maintains effective levels of the therapeutic diabody long-term, offering advantages over continuous diabody (blinatumomab) administration or CAR-T therapy, which comes with potential side effects (9).

A potency assay is essential for AAV vectors expressing CD3-CD19 diabodies to ensure consistent T-cell-mediated cytotoxicity against CD19-positive B cells, verifying the therapeutic’s biological activity before clinical use or batch release. This assay is critical for GMP compliance as it provides a quantifiable and reproducible measure of efficacy, ensures batch-to-batch consistency, and meets regulatory requirements for safety and effectiveness.

The goal was to develop a potency assay that quantifies the direct immune-mediated killing of tumor cells by CD3-CD19 diabodies. Existing cell-based reporter assays provide indirect measurements, which may not accurately capture T-cell-mediated cytotoxicity (9). There are reports that describe a direct cell-based immune killing assay using blinatumomab, in which PBMC-derived T cells were employed to evaluate tumor cell killing (10-13). However, PBMCs present significant challenges in developing and validating a GMP-compliant potency assay due to their high donor-to-donor variability, requirement for fresh isolation or cryopreservation, and inconsistent activation and expansion properties. These limitations complicate assay standardization, increase batch variability, and pose regulatory challenges for validation. For example, Li *et al*. (12) reported utilizing Nalm6, a human B-cell precursor leukemia cell line derived from a patient with acute lymphoblastic leukemia (ALL), in combination with human peripheral blood, Pan–T cells, and varying concentrations of blinatumomab. The assay exhibited low sensitivity, requiring more than 10 ng/mL of blinatumomab to achieve 50% killing of Nalm6 target tumor cells.

We describe a fluorescence-based potency assay that directly measures immune cell-mediated killing using the human cytotoxic T-cell line, TALL-104. In this method, RAJI cells (CD19-positive targets) are pre-labeled with calcein-AM dye, which remains contained within live cells. These target cells are co-cultured with the cytotoxic T cell line, TALL-104, and varying doses of GP101. Upon target cell death, the fluorescent dye is released, providing a quantifiable measure of cytotoxicity.

The method was validated to FDA GMP standards. The assay demonstrates excellent robustness, precision, and accuracy, and is suitable for measuring the potency of purified diabody, expressed from plasmid or AAV preparations, or samples from animal or human blood/serum. The method may be useful for validating other CD3-CD19 T cell engagers,

## 2. Materials and Methods

### 2.1. Reagents

The cytotoxic human T cell line, TALL-104 (CRL-11386) and B lymphoma cell line, RAJI (CCL-86), were obtained from the American Type Culture Collection (ATCC, Manassas,MD). Calcein-AM was obtained from Sigma (St, Louis, MO, USA). Fluorescence was measured using a Cytation 5 plate reader (Agilent, Santa Clara, CA). A Blinatumomab diabody was obtained from ProteoGenix (Schiltigheim, France). GP101 was produced by transduction of Expi293 cells with a plasmid containing the gene for GP101, then affinity purified on a column of CD19-coupled beads (ACROBiosystems, Newark, DE). The isolated protein was greater than 90% pure as seen by polyacrylamide gel electrophoresis, with the concentration determined by A280 using an extinction coefficient of 2.26 for 1 mg/mL based on amino acid content (4). This preparation is termed GP101 Reference Standard.

### 2.2. Cytotoxicity potency assay

A description of the cytotoxicity assay is shown in Fig. 1.

**Fig. 1.**
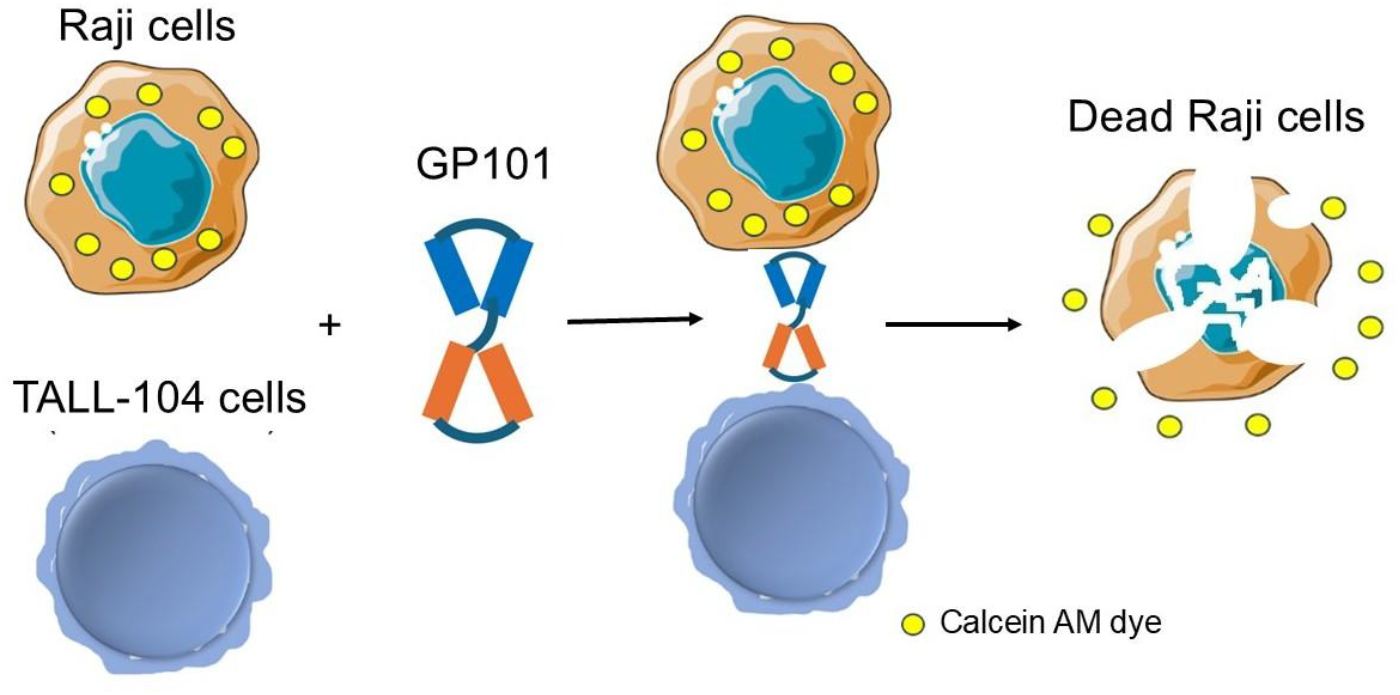
Schematic diagram of the cytotoxic potency assay. The RAJI target cells, labeled with a CalceinAM fluorescent dye, are incubated with cytotoxic TALL-104 lymphocytes and variable doses of bispecific GP101. After incubation, cells are centrifuged and the fluorescent dye released from dead cells is measured.

The TALL-104 cells were washed free of their Interleukin-2 (IL-2) growth factor and stored frozen in single use vials in liquid nitrogen. On the day of assay, cells were thawed, washed once, and 6,000 cells per well added to a 96 well plate in assay medium (RPMI1640, 10% Fetal Calf Serum, Penicillin/Streptomycin). The RAJI cells were incubated with 10uM CalceinAM for 1 hr at 37°C and washed to remove excellular CalceinAM. The dye-labeled RAJI, were added to the wells at a concentration of 2,000 cells per well. GP101 (partially purified) or Blinatumomab was added at increasing doses with a final volume of 0.2mL in the well. All diabody values shown in the data are final concentration in the wells. The 96 well plates were cultured at 37°C for 2 hr. All conditions were performed in triplicate or quadruplicate. The plates were centrifuged at 500 x g for 2 min and 100uL of the supernatant was removed to a black 96 well plate. Supernatant fluorescence was read in a BioTek Cytation.

### 2.3. GP101 stability studies using the cytotoxicity potency assay

The cytotoxicity potency assay is used to test the GP101 stability, which is evaluated by incubating the sample at a temperature of 2-8°C for a minimum duration of 2.5 weeks following thawing. Additionally, the stability study included an overnight incubation at 60°C, or 80°C. To assess potential changes over time, Reference Standard was compared to the samples stored at the various temperatures.

### 2.4. Statistical analysis

For potency results, the data were fit to a 4 Parameter (4P) curve using the Cytation instrument Gen5 software resulting in an EC_50_ value.

## 3. Results

### 3.1 Development of the method

Cytotoxic TALL-104 cells started killing dye-labeled RAJI targets within one hour (not shown) with > 90% killing after two hours of co-culture of the 2 cell types. The dye is stable in RAJI cells over this time frame but starts to be released when cells are cultured alone at later times (data not shown). Figure 2 shows a typical experiment, dilutions of Reference Standard fitted to a statistically determined 4P curve. The signal to noise ratio was less than 2.0; and the precision of replicates was within acceptable parameters. The results demonstrated a smooth dose response curve with an R^2^ fit of the data to the statistically generated curve is greater than 0.95, optimal for cell-based bioassays.

**Fig. 2.**
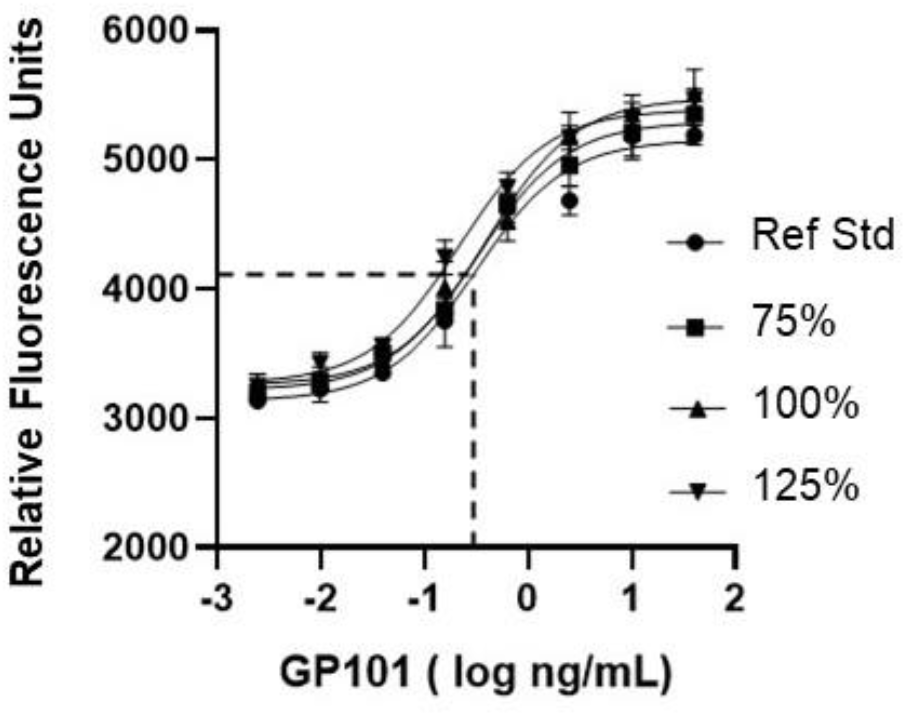
Assay of Reference Standard (100%) and of independent preparations at 70%, 100% and 130% of the 100% Reference Standard. Cytotoxic TALL-104 cells and Calcein AM dye-labeled RAJI target cells were co-cultured with GP101n for 2 hr. Toxicity was measured by release of fluorescent dye into the supernatant. Reference Standard (100%) and independent dilutions of Reference Standard at 70%, 100% and 130% were assayed in the same 96 well plate in triplicates. The graph shows 4P curve analysis of the data. The dotted lines indicate the 50% maximum fluorescence and the intercept on the x-axis is the EC_50_ value (ng/mL).

### 3.2. Linearity Accuracy of the method

The linearity of an analytical procedure is its ability to obtain test results which are directly proportional to the concentration of analyte. To test this, the Reference Standard was prepared at a 100%, termed as nominal, and independent dilutions were prepared at the same concentrations (100%) and also starting at 70% and 130%. Serial dilutions were prepared from these intermediate dilutions. These were tested in triplicate in one plate with the results in Figure 2. Analysis in Table 1 shows that the measured EC_50_ values for samples that were diluted from starting concentrations between 70% and 130% were within 4% of expected values. The plot of measured vs expected EC_50_ values should be a straight line with a slope of 1.00. For this experiment, the slope was 0.95 (Figure 3).

**Table 1.** 4P analysis for Reference Standard (Ref Std) and independent tests of Reference Standard at 100%, 70% and 130%.

| Test Name | Lower Plateau | Slope | EC50 ng/mL | Upper Plateau | R<sup>2</sup> | %Accuracy | %Agree Slope | %Agree Upper Plateau | %Agree Lower Plateau |
| --- | --- | --- | --- | --- | --- | --- | --- | --- | --- |
| 100% Ref Std | 3.09E+03 | 0.9 | 0.312 | 5.19E+03 | 0.969 |  |  |  |  |
| 100% Test | 3.19E+03 | 0.8 | 0.326 | 5.35E+03 | 0.979 | 96% | 93% | 103% | 103% |
| 70% Test% | 3.29E+03 | 0.9 | 0.434 | 5.53E+03 | 0.977 | 103% | 106% | 107% | 106% |
| 130% Test | 3.32E+03 | 0.9 | 0.242 | 5.42E+03 | 0.986 | 99% | 102% | 104% | 107% |

**Fig. 3.**
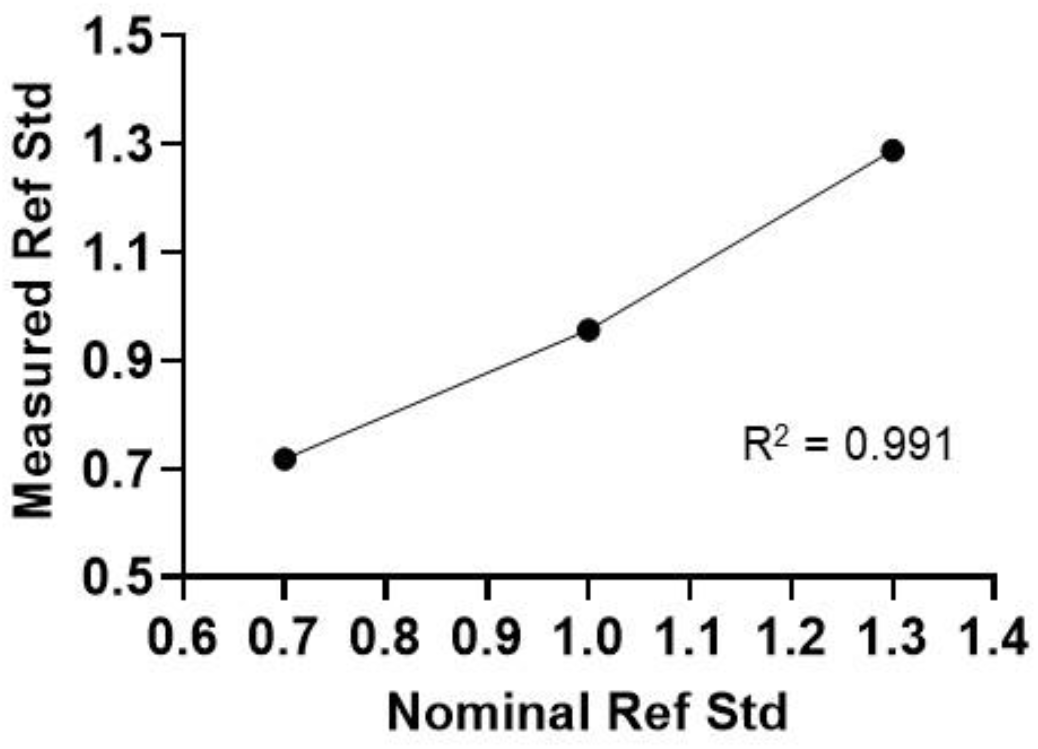
Plot of Measured vs Expected EC_50_ values for Reference Standard and tests of Reference Standard at 70%, 100% and 130%. Nominal is Reference Standard tested at dilutions starting at 40ng/mL.

### 3.3. Parallelism of a test sample to Reference Standard

To verify that a test sample is qualitatively similar to the Reference Standard, it should have similar biological effects. To measure this the Reference Standard and test sample curves should satisfy parallelism criteria. Table 2 shows that the statistically determined upper and lower plateaus (Relative Fluorescent Units) and slopes (no unit) agree within 93-107%.

**Table 2.** Parallelism test, 4P analysis for Reference Standard at 100% and independent tests of the Reference Standard at 100%, 70% and 130% of the nominal 100%.

| Test Name | Lower Plateau | Slope | EC50 ng/mL | Upper Plateau | R2 | %Accuracy | %Agree Slope | %Agree Upper Plateau | %Agree Lower Plateau |
| --- | --- | --- | --- | --- | --- | --- | --- | --- | --- |
| 100% Ref Std | 3.09E+03 | 0.9 | 0.312 | 5.19E+03 | 0.969 |  |  |  |  |
| 100% Test | 3.19E+03 | 0.8 | 0.326 | 5.35E+03 | 0.979 | 96% | 93% | 103% | 103% |
| 70% Test% | 3.29E+03 | 0.9 | 0.434 | 5.53E+03 | 0.977 | 103% | 106% | 107% | 106% |
| 130% Test | 3.32E+03 | 0.9 | 0.242 | 5.42E+03 | 0.986 | 99% | 102% | 104% | 107% |

### 3.4. Repeatability, the precision of triplicate fluorescent values

Triplicate fluorescent values at each Reference Standard and 100% test concentrations over 6 successive experiments had %CVs ranging from 0% to 6.0% (Table 3).

**Table 3.** Results of precision of triplicate values of Reference Standard and Test Samples, and EC_50_ values for Reference Standard.

| Experiment | %CV Range | Reference Standard EC50 ng/mL |
| --- | --- | --- |
| 1 | 0-5% | 0.626 |
| 2 | 1-7% | 0.651 |
| 3 | 1-6% | 0.304 |
| 4 | 0-6% | 0.224 |
| 5 | 0-5% | 0.324 |
| 6 | 0-5% | 0.374 |

### 3.5. Day-to-day precision

The precision of the assay to measure a test sample independently prepared reference Standard, compared to Reference Standard, over six successive experiments by two analysts are shown in Table 4. The data are shown as Relative Potency (RP) values, The EC_50_ of the test sample relative to that of Reference Standard. RP = [EC_50RE_ / [EC_50Test_]., a test sample weaker than Reference Standard needs more material to reach the E_50_ position. RP varied from 0.732 to 1.052 (Table 4).

**Table 4.** Results of intermediate precision. RP values on successive assay days by two analysts.

| Experiment | Analyst | RP of 100% sample |
| --- | --- | --- |
| 1 | 1 | 0.960 |
| 2 | 2 | 0.840 |
| 3 | 1 | 0.732 |
| 4 | 1 | 1.052 |
| 5 | 2 | 0.780 |
| 6 | 2 | 0.780 |
| Mean |  | 0.951 |
| %CV |  | 11.0 |
| Mean Analyst 1 |  | 0.915 |
| Mean Analyst 2 |  | 0.800 |

### 3.6. Ruggedness of the method

The robustness of the assay is acceptable. Effector and target cell concentrations can vary up to 20% with little effect on GP101 EC_50_ value; Calcein AM labeling concentrations can also vary up to 20% and give comparable results (not shown).

At a routine co-culture time of two hours, the EC_50_ for Reference Standard varies from about 0.2 to 0.5ng/mL in independent experiments; in the experiment shown it is 0.312ng/mL (Table 2). The R^2^ fit of the data to the statistically determined 4P curves are typically >0.95. Replicate but independent dilutions of GP101 assayed in the same 96 well plate show an accuracy of >90%. In Figure 2, test sample at 100% had an EC50 of 0.326ng/mL for an accuracy of 96%.

### 3.7. Stability of the assay

The Reference Standard EC_50_ value varies (Table 3), as expected for the response of cell lines on different days. We try to limit this variability by always using cells in log phase of cell growth and attention to other factors, such as short times out of the incubator, mild centrifugations. Note that both effector and target cells can vary from day to day and that the more important parameter of assay stability is the ability to measure RP, the potency of a test sample relative to Reference Standard. This parameter is tight, varying less than 20% from day to day.

### 3.8. Robustness

Robustness is shown by the comparable results of a second analyst, and by the precision of the RP values of a test sample from day to day (Table 4). Effector and target cell concentrations can vary up to 20% with little effect on GP101 EC_50_ value; Calcein AM labeling concentrations can also vary up to 20% and give comparable results (data not shown). The sensitivity of target RAJI cells to cytotoxicity was similar over eleven cell passages (data not shown).

### 3.9 Stability of GP-101 upon storage, and stability indicating property of the assay

Reference Standard has been stored at −80°C for up to one year with no loss in activity (not shown). Results show that GP-101 is stable for up to three Freeze /Thaws, and for at last 2 ½ weeks at 2 – 8°C. Incubation of GP101 overnight at 60°C and at 80°C totally destroyed activity (Table 5).

**Table 5.** GP101 Stability Data.

| Sample | EC50, ng/mL | Outcome |
| --- | --- | --- |
| RS | 0.487 |  |
| 2-8°C 2.5 weeks | 0.567 | Stable |
| 60°C O/N | >40 | Completely inactive |
| 80°C O/N | >40 | Completely inactive |
| Freeze/Thaw 3x | 0.433 | Stable |

### 3.10. Comparison of GP101 and blinatumomab

Blinatumomab shows a very similar potency to GP101, satisfying parallelism (top and bottom asymptotes within 20% and slope within 30%) and with a similar EC_50_. Figure 4 shows an example with EC_50_ of 1.16ng/mL for blinatumomab and 1.12ng/mL for GP101.

**Fig. 4.**
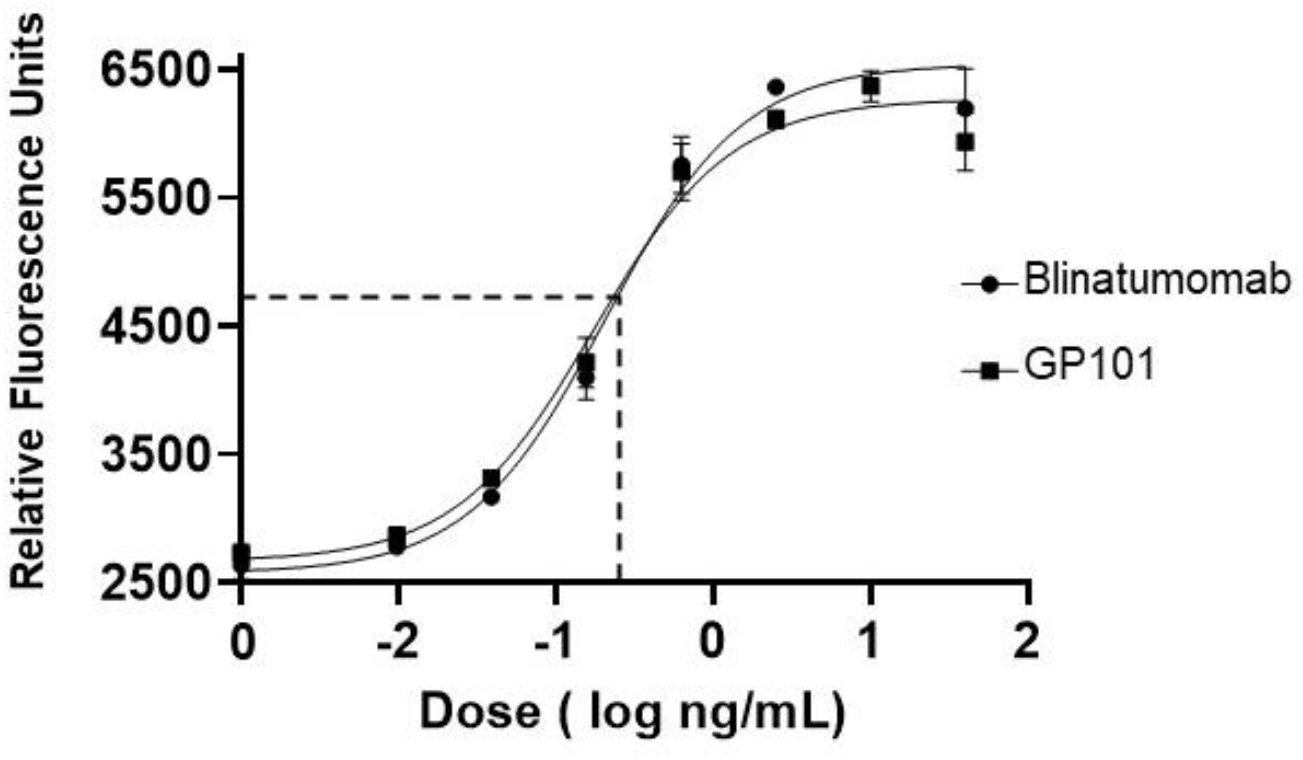
Cytotoxic assay of blinatumomab and GP101 in the same 96 well plate testing at the same concentration. The EC_50_ values, x-intercepts and the 50% maximum fluorescence, were 1.16 and 1.12ng/mL respectively.

### 3.11. Final assay acceptance criteria, System Suitability Criteria

For each experiment, the following criteria were created to demonstrate that the assay performed satisfactorily before reporting any EC_50_ or RP data (Table 6). The table shows the required criteria and the validation results.

**Table 6.**
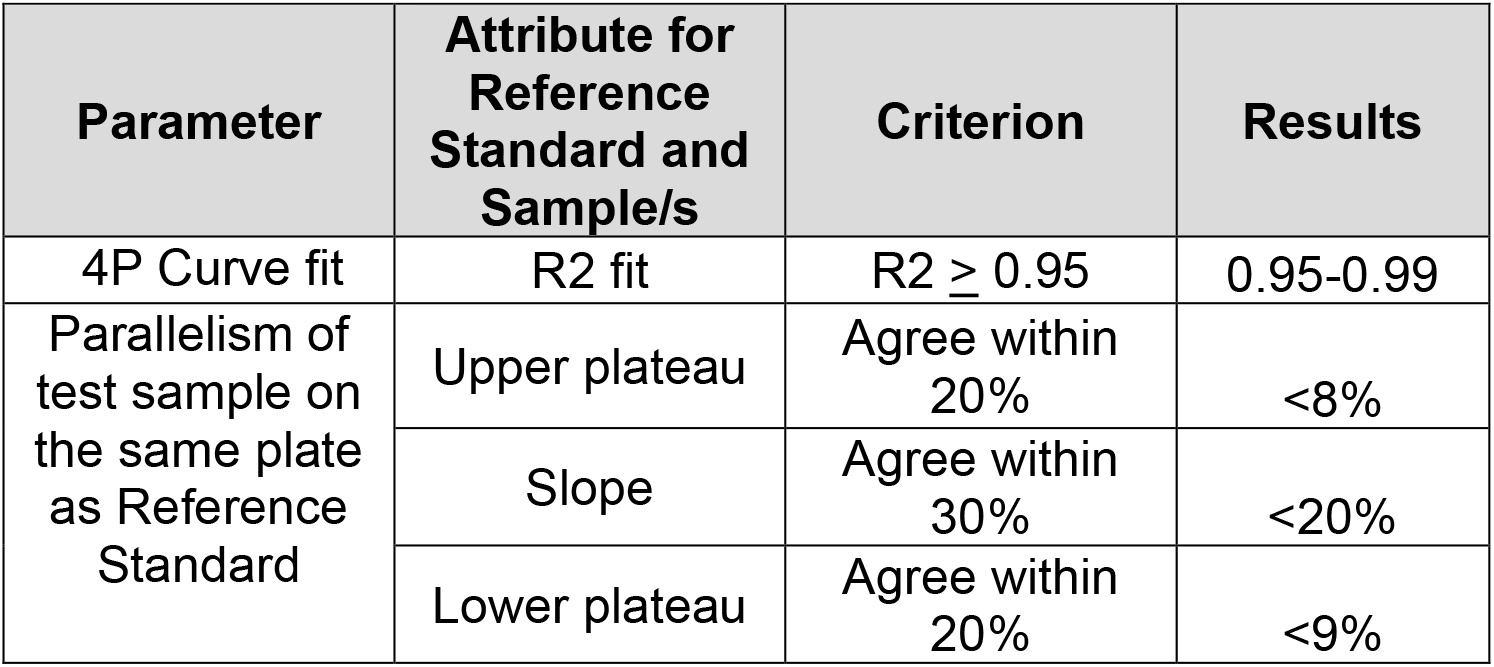
Assay Acceptance Criteria and Results.

| Parameter | Attribute for Reference Standard and Sample/s | Criterion | Results |
| --- | --- | --- | --- |
| 4P Curve fit | R2 fit | $R^2 \geq 0.95$ | 0.95-0.99 |
| Parallelism of test sample on the same plate as Reference Standard | Upper plateau | Agree within 20% | <8% |
|  | Slope | Agree within 30% | <20% |
|  | Lower plateau | Agree within 20% | <9% |

## 4. Discussion

The development of an Adeno-Associated Virus (AAV)-delivered diabody therapy, directing the GP101 diabody for treatment of B cell leukemias and lymphomas, represents a promising approach by enabling sustained therapeutic protein expression. However, ensuring consistent potency and quality of this therapeutic requires a robust and sensitive assay. The newly developed assay employs Calcein-AM dye-labeled RAJI cells as CD19-positive targets and CTLL-2 cells as cytotoxic effectors. The release of fluorescence upon cell death serves as a highly sensitive measure of GP101 activity. This assay demonstrated strong precision, accuracy, and robustness, with EC_50_ values consistently within the expected range. Furthermore, the method showed excellent repeatability, with independent dilutions of GP101 yielding accuracy greater than 90% and R^2^ values above 0.95 in 4P curve analyses. Parallelism testing confirmed that the potency curves of test samples closely matched those of the Reference Standard, reinforcing the assay’s reliability. Additionally, the fluorescence-based method exhibited strong linearity, with measured EC_50_ values deviating by less than 4% from expected values over a range of 70% to 130% of nominal 100% concentrations. The assay was also highly precise, with coefficient of variation (%CV) values below 7% across multiple experiments.

Robustness assessments demonstrated that slight variations in effector and target cell concentrations, as well as Calcein-AM labeling concentrations, had minimal impact on assay performance. Furthermore, the GP101 diabody displayed stability across multiple freeze/thaw cycles, storage at −80°C for over a year, and short-term stability at 2-8°C for up to 2.5 weeks. However, incubation at elevated temperatures (60°C and 80°C) led to complete loss of activity, confirming the stability-indicating capability of the assay.

Comparative testing with *blinatumomab*, a clinically approved bispecific T-cell engager, demonstrated similar potency, further validating GP101’s therapeutic relevance. The assay acceptance criteria established in this study provide a standardized framework to ensure the reliability of GP101 potency evaluations moving forward. Assays using this approach can be employed to measure diabody interaction between immune-mediated tumor cell killing expressing different antigens.

## Acknowledgments

We thank C.Murphy-Quin, and E.DiNello for experimentation and C. Fraser for assistance with the manuscript.

